# The structural proteome for the primordial glycolysis/gluconeogenesis

**DOI:** 10.1101/706192

**Authors:** Isabela Jerônimo Bezerra do Ó, Thais Gaudêncio Rego, Marco V. José, Sávio Torres de Farias

## Abstract

Comprehending the constitution of early biological metabolism is indispensable for the understanding of the origin and evolution of life on Earth. Here, we analyzed the structural proteome before the Last Universal Common Ancestor (LUCA) based in the reconstruction of the ancestral sequences and structure for proteins involved in glycolysis/gluconeogenesis. The results are compatible with the notion that the first portions of the proteins were the areas homologous to the present-day catalytic sites. Those “proto-proteins” had a simple function: binding to cofactors. Upon the accretion of new elements to the structure, the catalytic function could have emerged. Also, the first structural motifs might have been related to the emergence of the different proteins that work in modern organisms.

## Introduction

In “On the Origin of Species by Means of Natural Selection, or the Preservation of Favored Races in the Struggle for Life”, (1859) Charles Darwin suggested a common ancestor for all living organism. Since then the concept of the Last Universal Common Ancestor (LUCA), has triggered discussions about the components of this biological entity (Penny and Poole, 1999; Forterre et al., 2005; Mushegian, 2008; Glansdorff et al., 2008; Kim and Caetano-Anollés, 2011; Goldman et al., 2013).

One interpretation of the acronym LUCA refers to an already complex organism, with developed information processing, intricate metabolism and a DNA-based genome (Di Giulio, 2003). Thus, LUCA does not represent the earliest form of life, because the evolutionary processes had begun long before DNA. Carl Woese (1998) suggested the existence of communities of progenotes as precursors to LUCA. Progenotes were membrane-delimited organisms with few metabolic functions. In this scenario, these organisms or biological entities had RNA as informational molecule, and an imprecise translation process, generating statistical proteins, i.e., sequences of peptides arranged probabilistically, which would be under the influence of natural selection. We can describe the progenotes as a community of subsystems that coevolved in a way that at a particular time of development it was established a higher stage of dependence among them, which culminated in the congregation of these subsystems (Woese, 1998).

It stands to reason to assume that not all steps of the modern metabolic pathways were present in a progenote; different steps of metabolism appeared in different progenotes. The ability to translate certain types of information was most likely one of the first properties, if not the first, to develop among progenotes (Eigen and Schuster, 1978; Woese, 1998).

According to Weiss et al. (2016), there were traces of carbon, nitrogen and energy metabolism in the genome of the LUCA. The carbon metabolism was most likely present in the earliest forms of life as we know it today, arguably not only by its vast presence in all life domains but also by its importance in generating energy and precursors for biological systems. From the carbohydrate pathways, the two most universal and well preserved are glycolysis/gluconeogenesis and citric acid cycle.

The antiquity of these pathways, specifically gycolysis/Gluconeogenesis, is suggested by many features. First, glycolysis/gluconeogenesis is present, at least partially in all life forms, and all cellular life carry variations of carbohydrate metabolism. Second, their structural simplicity, meaning that the enzymes act independently and do not require membranes, which would be a necessary characteristic in an Archean ocean on a pre-membrane reality. Thirdly, it is independent of oxygen, another feature necessary at that time.

Additionally, the results of the two substrate-level phosphorylation reactions would provide enough energy for primitive metabolism. With pyruvate as the terminal electron acceptor, the redox balance of the pathway can be maintained. Another reason to sustain this pathway being prior to LUCA is that glycolysis evolved most likely as a reverse Gluconeogenesis. The synthesis of hexose sugars would act as a storage of energy and biological precursors, which would be advantageous in a primitive scenario with scarce complex molecules.

At this point in evolutionary history, large accumulation of sugar was not feasible. Glycolysis is not only responsible for energy output but its products three-carbon compounds (eg dihydroxyacetone phosphate and glyceraldehyde-3-phosphate) act as a source of energy as well. These products are used in a number of other pathways such as the amino acid anabolism. Producing the three-carbon compounds was probably the first role of glycolysis, which is supported by it being constant in all life forms while the energy metabolism varies. Storing carbon was probably advantageous as well, as it decreased organisms’ dependence of the system surrounding it.

Large informational molecules most probably did not have sufficient stability on primitive Earth, and the first coding devices must have reflected the chemical constraints of their origin. Eigen and Winkler-Oswatitsch (1981) proposed that the initial molecular events were carried out by tRNAs, mostly because: (1) they have predominant GC pairing, necessary for relative stability within lengths of tRNAs; (2) they have stable folded secondary structure, enhancing their resistance to hydrolytic decomposition; (3) the codons act as messenger carriers; (4) it has around 50 to 100 residues ideal for prebiotic polymerization conditions; (5) ancient family tree; and (6) symmetric structure.

Eigen and Schuster (1978) suggested that the first anticodons were of the type RNY (purine-any nucleotide-pyrimidine). These are codons where the first base is a purine, the third a pyrimidine, and the second is any of them. Hence, the first modules were rich in alanine, serine, threonine, asparagine, aspartic acid, valine, isoleucine, and glycine.

Farias et al (2016) reconstructed the ancestral sequence of tRNAs that may have acted as the first genes of these progenotes. They proposed a proteome for the progenotes based on the translation of ancestral tRNAs with a RNY pattern. Among the various metabolic pathways found in modern organisms, some were already present in LUCA and on the progenotes. Farias et al (2016) by analyzing the similarities of translated tRNAs with modern proteins, found possible homologies with the following pathways: Amino acids pathways, glycolysis/gluconeogenesis, lipids pathways, nucleotides pathways, translation, and transcription or RNA replication.

Here we use the data obtained by Farias et al. (2016) and their suggestions for the early metabolism to reconstruct two different stages of evolution. One (bottom up) prior to the Last Universal Common Ancestor which will be based on the proteome established by tRNA ancestor molecules and another (top down) based in the reconstruction of ancestor sequence from modern proteins involved in the metabolism of Glycolysis/Gluconeogenesis and three carbons compound and compare both strategies to understand the evolutionary history of this metabolic pathway.

## Material and Methods

### Ancestral Approaches

Two different approaches were used to reconstruct ancestors.

### Bottom Up approach

The first one, “bottom up”, refers to the reconstructions based on the sequences found by Farias et al. (2016a – supplementary information). The proteome suggested by them was made using sequences of ancestral tRNAs following the RNY pattern, that when translated presented matches with modern proteins involved in the three carbons metabolism.

### Top Down approach

The other approach was a reconstruction using sequences of the modern proteins chosen based on the ancestral proteome suggested by Farias et al. (2016b) which were the following: glycerate kinase, triosephosphate isomerase, glucose 6-phosphate 1-dehydrogenase 2, glucose 6-phosphate isomerase, phosphoglycerate kinase, glycerol 3-phosphate dehydrogenase and transketolase.

### Ancestral sequence reconstruction

We obtained 400 sequences for each type of enzymes that are part of the glycolysis/gluconeogenesis and three carbons metabolic pathways from the National Center for Biotechnology Information (NCBI) (supplementary information). The sequences covered the three domains of life, namely Archaea, Bacteria and Eukarya. The sequences were aligned with the support of the software MAFFT v.7 (Katoh and Standley, 2016). To align the sequences, we used BLOSUM45 for the scoring matrix, and all the other parameters remained as default. To reconstruct the most probable ancestral sequence, we used maximum likelihood with the support of software MEGA7 (Kumar et al., 2015). For this approach, we used different levels of strictness for inferring the ancestor sequence by maximum likelihood, were used complete deletion, 95% and 99% site coverage cutoff.

### Structural analysis

The proteins structures were reconstructed by homology using the web server I-Tasser (http://zhanglab.ccmb.med.umich.edu/I-TASSER/) (Zhang, 2008). The templates of modern proteins and the ligands were obtained in Protein Data Base (www.rcsb.org) (Berman et al., 2000).

The structural alignments were performed using the software TM-Align (http://zhanglab.ccmb.med.umich.edu/TM-align/)(Zhang and Skolnick, 2005). For analysis of the ligands to the ancestral structure in both approaches (top-down and bottom-up), we used the software Coach (http://zhanglab.ccmb.med.umich.edu/COACH/) (Yang et al., 2013).

### Results and Discussion

In the center of the modern metabolism are the glycolysis/gluconeogenesis pathways, which are conserved in all domains of life. These pathways are the core of the carbon metabolism in modern cells. It has been suggested that glycolysis may have originated as a mechanism that extracted energy and biological precursors from abiotically formed amino acids (Kenneth et al., 2004). Sutherland (2017) suggested the importance of the compounds with three carbons in the emergence of the first metabolic pathways. Farias et al. (2016) proposed that these compounds worked as a distribution center of carbon to other metabolic pathways in development (Jheeta, 2015).

Farias et al. (2016) reconstructed the ancestral sequence from tRNAs and compared the ancestor sequences translated with the modern proteins. The results suggested that the enzymes of glycolysis/gluconeogenesis metabolism could have emerged from proto-tRNAs as well as the first genes, the proteins involved in the metabolism of three carbon compounds being the core of the process. Here, we call this approach bottom up, where the structures are derived from translated sequences of the tRNA ancestor.

To understand the structural evolutionary history of the Glycolysis/Gluconeogenesis enzymes, we used a second approach based in the reconstruction of the ancestral sequences and structures from the modern enzymes involved in Glycolysis/Gluconeogenesis and three carbon compound (top down approach). We used just the same enzymes that had matches with translated sequences of the tRNA ancestor (see materials and methods), to make comparisons between the structures obtained from different approaches. From the ancestral sequences obtained from modern enzymes, we used complete deletion, 95% and 99% site coverage cutoff, which may capture similarities with different steps in the evolutionary history of these enzymes.

### Ancestral structures reconstruction

Figure 1, the reconstructed structures by homology of the tRNA ancestor sequences translated in the bottom up approach. In general, the structures are composed by simple motifs, basically, loops and small alpha helices. Table 1 shows the most similar modern structures with RMSD values (root-mean-square deviation) between ancestor by bottom up approach and modern proteins, and the organisms from which the protein belongs.

**Figure 1.**
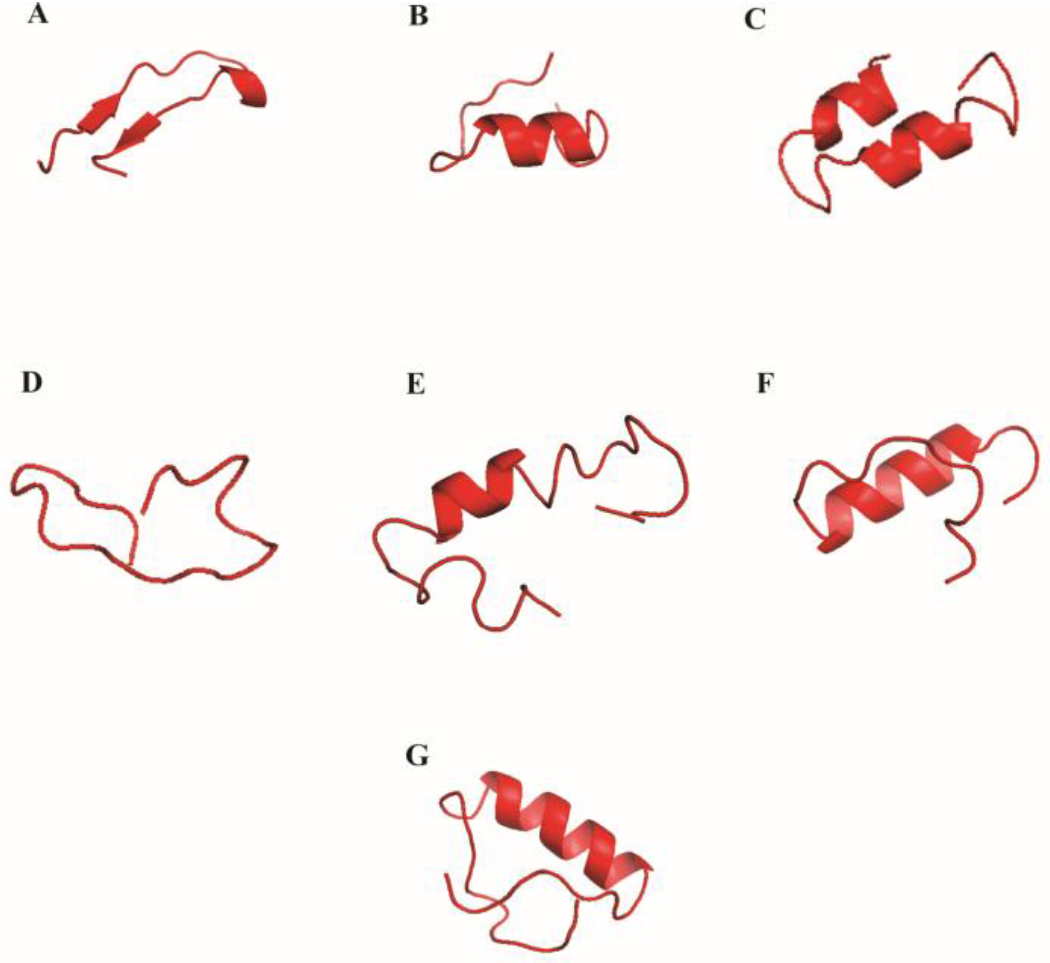
Structures of ancestral sequences obtained by the bottom up approach. A) glyceraldehyde-3-phosphate-dehydrogenase, B) glucose 6-phosphate1-dehydrogenase, C) glucose 6 Phosphate Isomerase, D) glycerate Kinase, E) phosphoglycerate kinase, F) triose phosphate isomerase and G) transketolase

**Table 1.** Ancestral proteins obtained from bottom up approach, the best similar structure in modern organisms and the RMSD distance between the ancestral protein and modern protein.

| Bottom Up | Most Similar Structure | Organism | RMSD |
| --- | --- | --- | --- |
| **Glucose-6-phosphate Isomerase** | Iron ABC transporter solute-binding protein (4HN9) | *Eubacterium eligens* | 1.69 |
| **Phosphoglycerate-kinase** | RNA binding domain (4GH9) | *Marburg virus* | 2.67 |
| **Triosephosphate-isomerase** | Hydrolase (Crystal structure of had family enzyme bt-2542 target e fi-501088) magnesium complex (4DFD) | *Bacteroides thetaiotaomicron* | 1.46 |
| **Glycerate-kinase** | Kinase domain of protein tyrosine kinase 2 beta (3CC6) | *Homo sapiens* | 1,95 |
| **Glyceraldehyde-3-phosphate-dehydrogenase** | Enolase superfamily member complexed with Mg and Glycerol (3R25) | *Vibrionales bacterium* | 2.10 |
| **Glucose-6-phosphate-1-dehydrogenase** | CRISPR-associated protein Cas6e (4DZD) | *Escherichia coli* | 1.78 |
| **Transketolase** | Acyl-CoA dehydrogenase fold (5AHS) | *Advenella mimigardefordensis* | 1.99 |

These results are in agreement with basic functions for very early proteins in the beginning of the biological systems, when sophisticated catalytic functions were evolving. The initial proteins were very simple and were able to bind to a variety of substrates as they were rudimentary and had low affinity – work of Frances Arnold, 2018 Nobel Prize Laureate in chemistry (Newton, 2018). The similarities with proteins that are not involved in carbon metabolism can indicate that these basic motifs worked as mounting blocks in the origin of the first proteins and genes.

The most similar modern structures showed basic biological functions such as hydrolase, RNA binding domain, enolase, among other, with a range of RMSD value of 1.46 to 2.67. These results can indicate the basic structural blocks that participated in the formation of many different pathways, and the combination of these structural parts was relevant to primordial diversification of essential pathways and for the establishment and maintenance of the emerging biological systems.

Figure 2 shows the structures constructed by homology by the top down approach with complete deletion and 99% site coverage cutoff and in Figure 3 with 95% site coverage cutoff. The comparison between the structures obtained from the top down and the bottom up approaches indicates that the structures from the top down approach exhibit a more complex structure.

**Figure 2.**
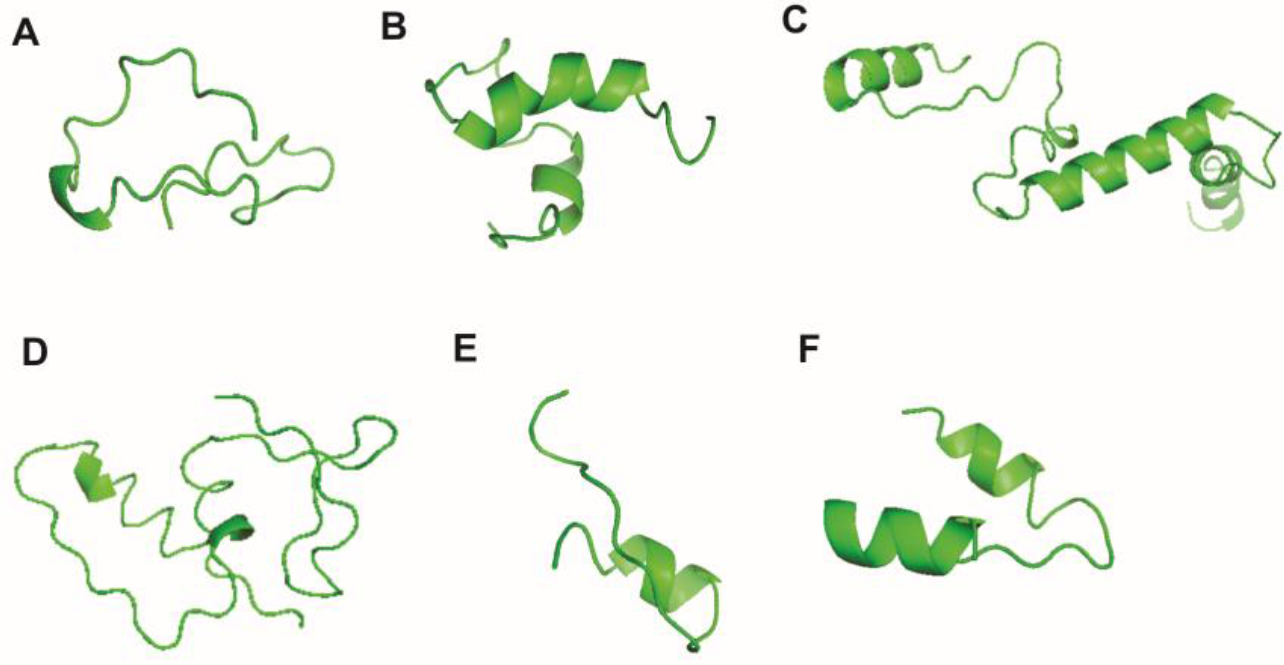
Structures to ancestral sequences obtained by top down approach with complete deletion (CD) and 99% site coverage cutoff (PD99). A) glyceraldehyde-3-phosphate-dehydrogenase (PD99), B) glucose 6-phosphate1-dehydrogenase (CD), C) glucose 6 phosphate Isomerase (PD99), D) glycerate Kinase (CD), (E) triose phosphate isomerase (CD) and (F) transketolase (PD99).

**Figure 3.**
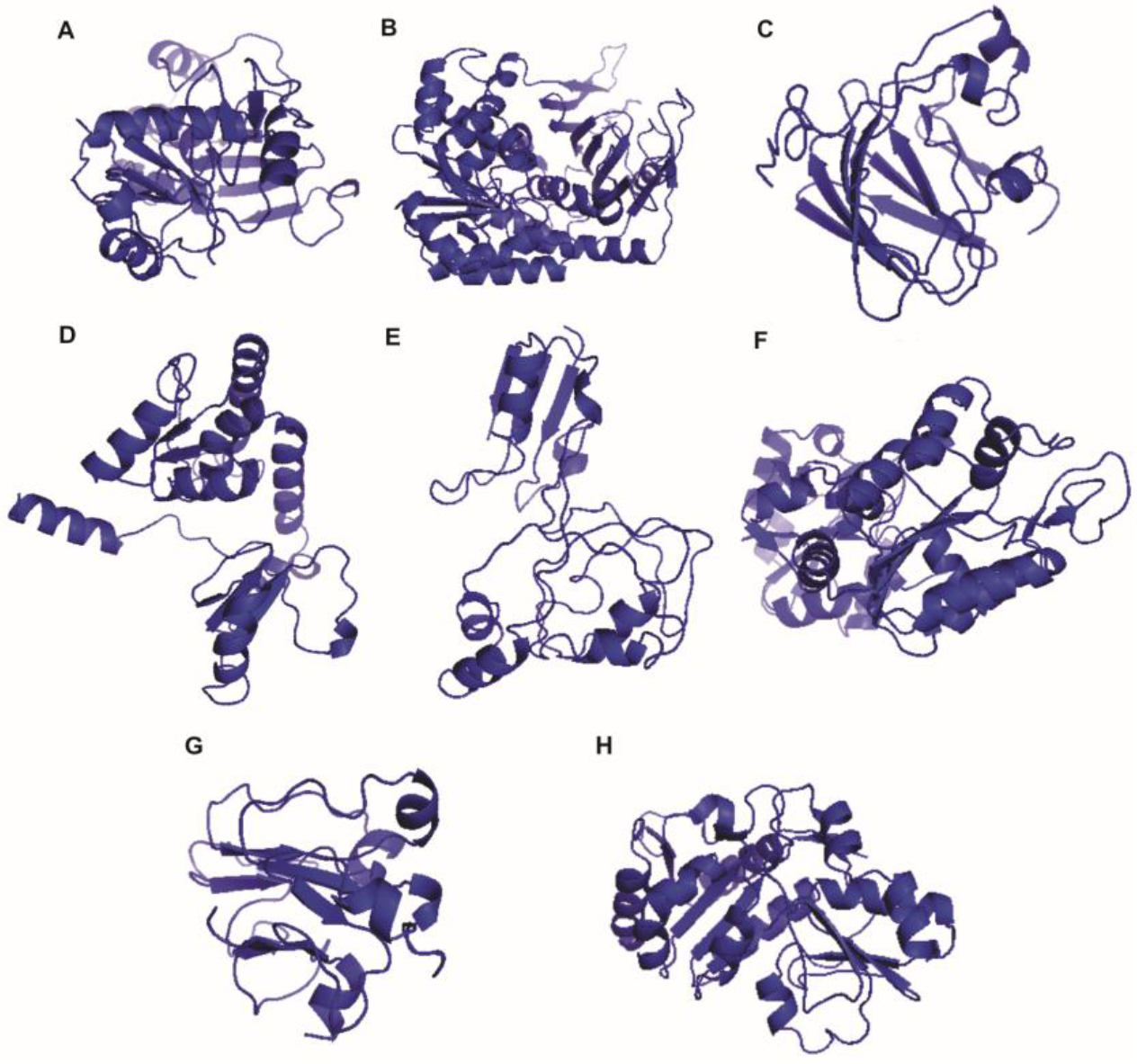
Structures to ancestral sequences obtained by top down approach with 95% site coverage cutoff. A) glyceraldehyde-3-phosphate-dehydrogenase, B) glucose 6-phosphate1-dehydrogenase, C) glucose 6 phosphate Isomerase (I), D) glucose 6 phosphate isomerase (II), E) glycerate kinase, F) phosphoglycerate kinase, G) triose phosphate isomerase and H) transketolase.

In Table 2, the most similar modern structures with RMSD values between ancestor by top down approach and modern proteins are shown. These results reflect the limitations of the method to reconstruct ancestral sequences from modern proteins, where it is possible to identify the first functional domains, but not the deep evolution of these domains, before the establishment of the modern specificity. However, these results also show the efficiency of the method to understand the evolutionary history of proteins until the first functional domain.

**Table 2.**
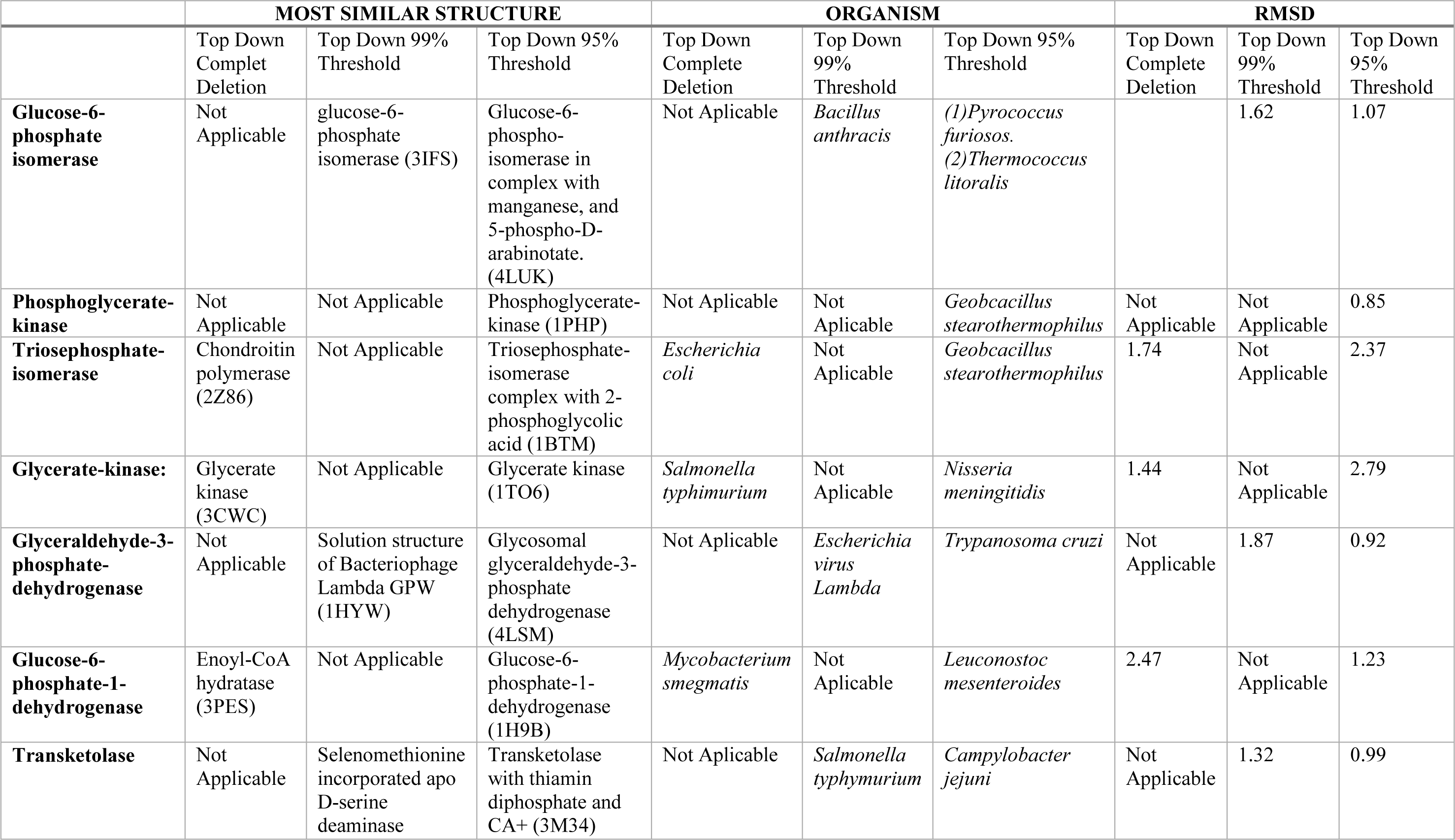
Ancestral proteins obtained from Top down approach, the best similar structure in modern organisms and the RMSD distance between the ancestral protein and modern protein. In parentheses, the protein database code.

|  | MOST SIMILAR STRUCTURE |  |  | ORGANISM |  |  | RMSD |  |  |
| --- | --- | --- | --- | --- | --- | --- | --- | --- | --- |
|  | Top Down<br>Comple<br>Deletion | Top Down 99%<br>Threshold | Top Down 95%<br>Threshold | Top Down<br>Complete<br>Deletion | Top Down<br>99%<br>Threshold | Top Down 95%<br>Threshold | Top Down<br>Complete<br>Deletion | Top Down<br>99%<br>Threshold | Top Down<br>95%<br>Threshold |
| **Glucose-6-phosphate isomerase** | Not Applicable | glucose-6-phosphate isomerase (3IFS) | Glucose-6-phospho-isomerase in complex with manganese, and 5-phospho-D-arabinotate. (4LUK) | Not Aplicable | *Bacillus anthracis* | (1)*Pyrococcus furiosus*.<br>(2)*Thermococcus litoralis* |  | 1.62 | 1.07 |
| **Phosphoglycerate-kinase** | Not Applicable | Not Applicable | Phosphoglycerate-kinase (1PHP) | Not Aplicable | Not Applicable | *Geobacillus stearothermophilus* | Not Applicable | Not Applicable | 0.85 |
| **Triosephosphate-isomerase** | Chondroitin polymerase (2Z86) | Not Applicable | Triosephosphate-isomerase complex with 2-phosphoglycolic acid (1BTM) | *Escherichia coli* | Not Applicable | *Geobacillus stearothermophilus* | 1.74 | Not Applicable | 2.37 |
| **Glycerate-kinase:** | Glycerate kinase (3CWC) | Not Applicable | Glycerate kinase (1TO6) | *Salmonella typhimurium* | Not Applicable | *Nisseria meningitidis* | 1.44 | Not Applicable | 2.79 |
| **Glyceraldehyde-3-phosphate-dehydrogenase** | Not Applicable | Solution structure of Bacteriophage Lambda GPW (1HYW) | Glycosomal glyceraldehyde-3-phosphate dehydrogenase (4LSM) | Not Applicable | *Escherichia virus Lambda* | *Trypanosoma cruzi* | Not Applicable | 1.87 | 0.92 |
| **Glucose-6-phosphate-1-dehydrogenase** | Enoyl-CoA hydratase (3PES) | Not Applicable | Glucose-6-phosphate-1-dehydrogenase (1H9B) | *Mycobacterium smegmatis* | Not Applicable | *Leuconostoc mesenteroides* | 2.47 | Not Applicable | 1.23 |
| **Transketolase** | Not Applicable | Selenomethionine incorporated apo D-serine deaminase | Transketolase with thiamin diphosphate and CA+ (3M34) | Not Applicable | *Salmonella typhimurium* | *Campylobacter jejuni* | Not Applicable | 1.32 | 0.99 |

Our results show that the suggested ancestral structures are similar to structures of different extremophiles that lived in either acidic habitats or in high temperatures, or both. This may indicate that these first enzymes before and after LUCA were adapted to extreme scenarios to reflect the environmental conditions on the Earth some 4.0 billion years ago (Bell et al, 2015, Jheeta, 2017). It is suggested that Theia’s impact with the Earth created enough energy to melt Earth’s mantle. The condensed water formed a dense atmosphere obstructing the dissipation of heat which resulted in a greenhouse effect (Nisbet et al., 2007). Even though there is still a debate, different pieces of evidence seem to show that the Earth might have reached extremely high temperatures on the hadean eon and this might have influenced biological evolution (Sleep et al., 2007; Zahnle et al., 2007).

### Analysis of substrates binding

Table 3 shows the analysis of substrate binding for the ancestral structures obtained for the bottom up and the top down approaches. For the bottom up approach, the results of substrate binding indicate that these structures could interact with simple compounds that may have worked as cofactors for basic functions. Some of the structures bind to amino acids, most of them bind to coenzymes that have the potential of energy transfer, and all of them bind to small hydrocarbons. The interaction between the ancestral structures and nucleotides also suggest that the coevolutionary relationship between these compounds were established earlier in the primordial environment.

**Table 3.**
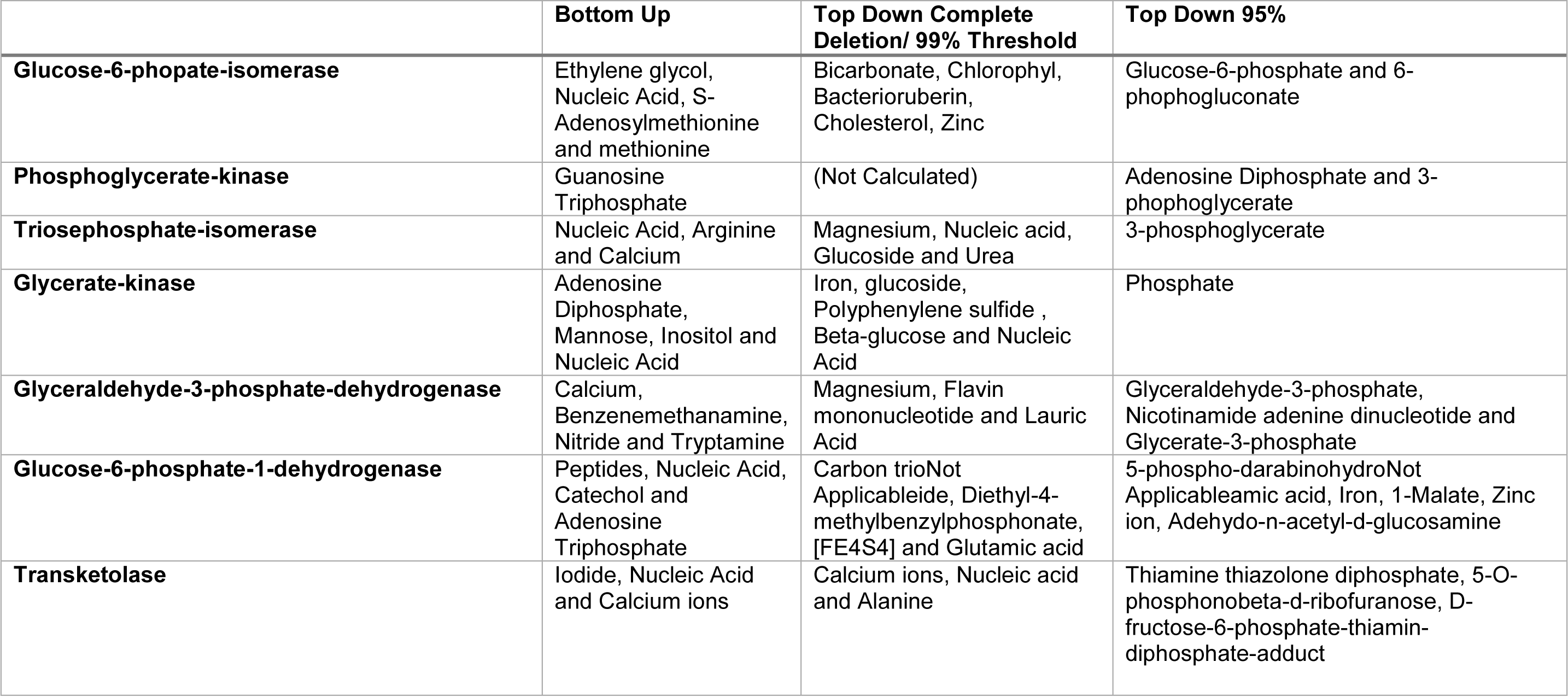
The ancestral structures obtained by Bottom up and top down approach and the best ligands for each ancestral protein.

Caetano-Anolles et al. (2012) suggested that in the origin of the initial metabolic pathways, the progressive appearance of complexity was the rule and that the first proteins or peptides worked by binding various cofactors, and the accretion of new parts made it possible for these primitive proteins to be able to operate on catalysis and molecular functions. As in modern biological systems, the model suggests that nucleic acids worked as cofactors in the beginnings of life. For the structures obtained by the top down approach with complete deletion and 99% site coverage cutoff, we observed a similar pattern of substrate binding with the structures obtained from bottom up approach. For the structures obtained by the top down approach with 95% site coverage cutoff, the best substrates were similar to the modern proteins. These results suggest that during the evolutionary history, the accretion of new structural parts in the protein made the structure more complex and specific. Increased specificity, enabled the fixation of the first metabolic pathways.

### Evolutionary steps in the natural history of Glycolysis/Gluconeogenesis

In Figure 4, the evolutionary steps are reconstructed. The data suggests that the initial portion of the proteins were simple structural motifs near the homologous are to the modern catalytic site. During the evolutionary accretion of new motifs, selective pressure worked to protect and isolate the catalytic site, making the protein more specific. Alves et al. (2002) showed that functional blocks of similar chemistry evolved within the metabolic networks. The bottom up ancestors were all structurally close, more than they were to their homologs. These ancestors also had similar substrates but bound to a variety of ligands. This could indicate that they had a common moiety that had a non-specific action. These first “enzyme-like” molecules could have acted at different points of the metabolism, and throughout evolution they gained specificity. Other than gaining specificity, the structure of the enzymes became larger and more complex. The evolution probably happened around the catalytic site following the accretion model.

**Figure 4.**
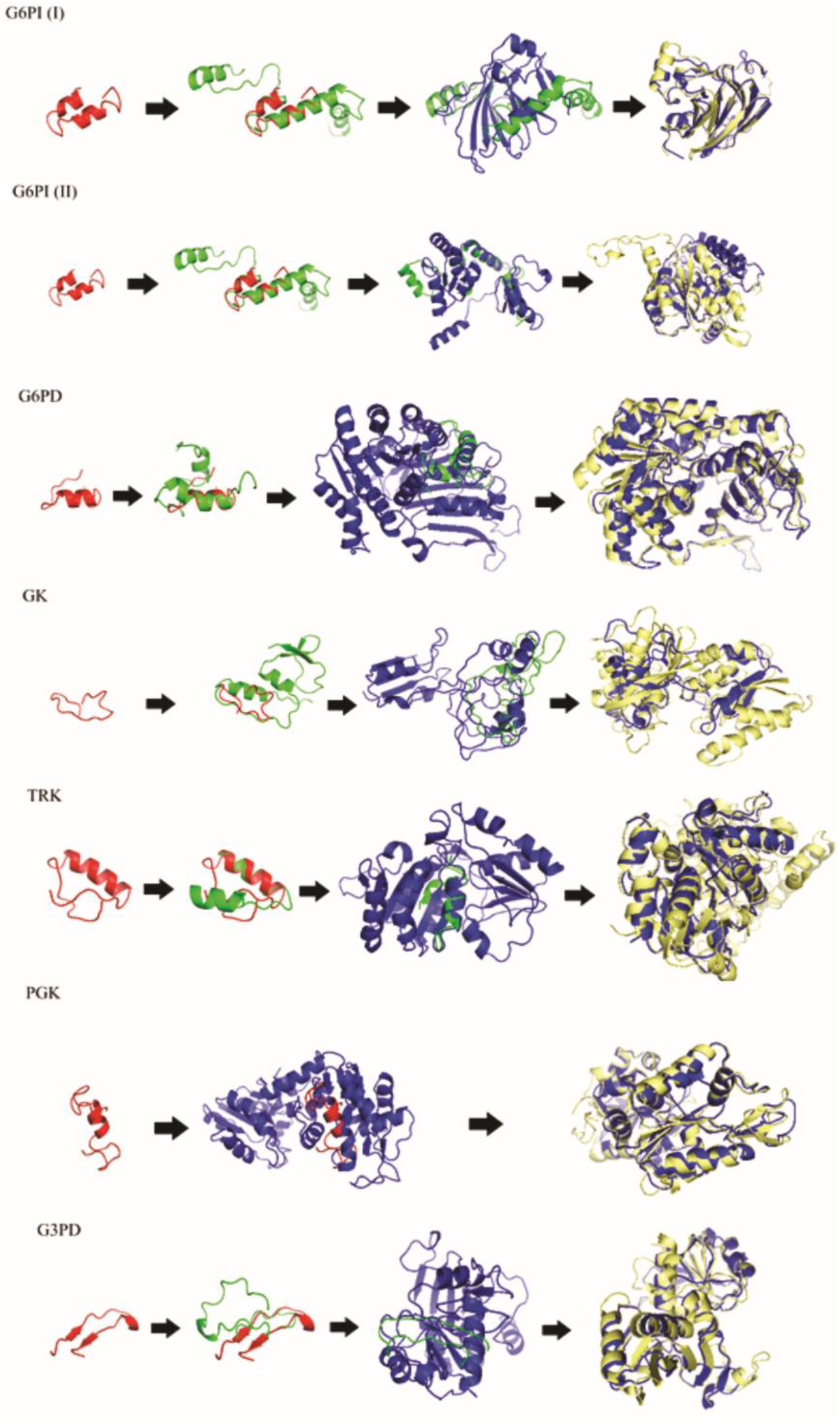
Proposed evolutionary steps of structures. Alignment made using TM-Align server. In red the bottom up ancestors, in green the top down ancestors calculated with complete deletion or the top down ancestors calculated with partial deletion with 99% threshold, in blue the top down ancestors calculated with 95% threshold and in pale yellow the modern enzymes. gyceraldehyde-3-phosphate-dehydrogenase (G3PD), glucose 6-phosphate1-dehydrogenase G6PD, glucose 6 phosphate isomerase (G6PI (I) and (II)), glycerate kinase (GK), phosphoglycerate kinase (PGK), triose phosphate isomerase (TPI) and transketolase (TRK).

We contend that the first action of these enzymes related to glycolysis was actually in the gluconeogenic direction. The synthesis of hydrocarbons would be positively selected since it provided both stored energy and biosynthetic precursors.

## Conclusion

The comprehension of the origin and evolution of the metabolic pathways is a keystone to the understanding of the emergence of life itself. Here, we analyzed the origin and evolution of the proteins involved in glycolysis/gluconeogenesis. The results endorse that the initial genetic information may have originated from proto-tRNAs. As well as that, the first peptides worked simply by binding cofactors. Accretion of new motifs resulted in complexification of the proteic structure, which allowed for the sophistication of the catalytic domain.

Our work indicates that in the emergence of the first peptides, simple structural motifs worked as mounting blocks and by the combination of the different motifs, different proteins emerged that work in modern organisms. The evolution of the proteins involved in glycolysis/gluconeogenesis occurred by selective pressure to protect the catalytic domain, which culminated in the improvement of the catalysis. We suggest that gluconeogenesis preceded glycolysis and, with the accumulation of the carbon in the initial biological system, a balance between the pathways was obtained.

## References

Alves, R., Chaleil, R.A., Sternberg, M.J., 2002. Evolution of enzymes in metabolism: a network perspective. J. Mol. Biol. 19;320(4):751–70.

Bell, E.A., Boehnke, P., Harrison, T.M. and Mao, W.L. 2015. Potentially biogenic carbon preserved in a 4.1 billion-year-old zircon. Proceedings of the National Academy of Sciences of the United States of America. 112(47), 14518–14521

Berman, H.M., Westbrook, J., Feng, Z., Gilliland, G., Bhat, T.N., Weissig, H., Shindyalov, I.N., Bourne, P.E., 2000. The Protein Data Bank. Nucleic Acids Res. 1;28(1):235–42.

Caetano-Anollés, G., Kim, K.M., Caetano-Anollés, D., 2012. The phylogenomic roots of modern biochemistry: origins of proteins, cofactors and protein biosynthesis. J. Mol. Evol. 74(1-2):1–34. doi: 10.1007/s00239-011-9480-1.

Di Giulio, M., 2003. The universal ancestor and the ancestor of bacteria were hyperthermophiles. J. Mol. Evol. 57(6):721–30.

Eigen, M., Schuster, P., 1978. The hypercycle. A principle of natural self-organization. Part A: Emergence of the hypercycle. Naturwissenschaften, 64(11):541–65.

Eigen, M., Winkler-Oswatitsch, R., 1981. Transfer-RNA, an early gene? Naturwissenschaften, 68(6):282–92.

Farias, S.T., Rêgo, T.G., José, M.V. 2016a. tRNA Core Hypothesis for the Transition from the RNA World to the Ribonucleoprotein World. Life (Basel). 23;6(2). pii: E15. doi: 10.3390/life6020015.

Farias, S.T, Rego, T.G., Jose, M.V., 2016b. A proposal of the proteome before the last universal common ancestor (LUCA). Int. J. Astrobiol. 15 (1): 27–31. doi:10.1017/S1473550415000464.

Forterre, P., Gribaldo, S., Brochier, C., 2005. Luca: the last universal common ancestor. Med. Sci. (Paris). 21(10):860–5.

Glansdorff, N., Xu, Y., Labedan, B., 2008. The last universal common ancestor: emergence, constitution and genetic legacy of an elusive forerunner. Biol. Direct, 9;3:29. doi: 10.1186/1745-6150-3-29.

Goldman, A.D., Bernhard, T.M., Dolzhenko, E., Landweber, L.F. 2013. LUCApedia: a database for the study of ancient life. Nucleic Acids Res. 41(Database issue):D1079–82. doi: 10.1093/nar/gks1217.

Katoh, K., Standley, D.M., 2016. A simple method to control over-alignment in the MAFFT multiple sequence alignment program. Bioinformatics, 1;32(13):1933–42. doi: 10.1093/bioinformatics/btw108.

Kenneth, S.B. 2004. Functional Metabolism: Regulation and Adaptation. John Wiley & Sons, INC., Publication.

Kim, K.M., Caetano-Anollés, g., 2011. The proteomic complexity and rise of the primordial ancestor of diversified life. BMC Evol. Biol. 25;11:140. doi:10.1186/1471-2148-11-140.

Kumar, S., Stecher, G., Tamura, K., 2016. MEGA7: Moelcular evolutionary genetics analysis version 7.0 for bigger datasets. Mol. Biol. Evol. 33 (7): 1870–1874. doi:10.1093/molbev/msw054.

Jheeta, S. 2015. Conference report: the Routes of emergence of life from LUCA during the RNA and viral world: a conspectus. Life, 5, 1445–1453; doi:10.3390/life5021445.

Jheeta, S., 2017. Conference Report The Landscape of the Emergence of Life. Life, 7, 27; doi:10.3390/life7020027.

Mushegian, A., 2008. Gene content of LUCA, the last universal common ancestor. Front. Biosci. 1;13:4657–66.

Newton, J., (2018). The potential of promiscuity. Chemistry World, 15(11), 28–29

Nisbet, E., Zahnle, K., Gerasimov, M.V, Helbert, J., Jaumann, R., Hofmann, B.A,, Benzerara, K., Westall, F., 2007. Creating habitable zones, at all scales, from planets to mud micro-habitats, on Earth and on Mars. Space Sci. Rev. 129:79–121.

Penny, D., Poole, A., 1999. The nature of the last universal common ancestor. Curr. Opin. Genetics Dev. 9(6):672–7.

Sleep, N.H. 2007. Plate tectonics through time, Treatise on Geophysics Volume 9, (ed. Schubert G.), pp. 101–117 Oxford, Elsevier.

Sutherland, J.D. 2017. Studies on the origin of life – the end of the beginning. Nat. Rev. Chem. 01:12. DOI/10.1038/s41570-016-0012.

Weiss, M.C., Sousa, F.L., Mrnjavac, N., Neukirchen, S., Roettger, M., Nelson-Sathi, S., Martin, W.F. 2016. The physiology and habitat of the last universal common ancestor. Nat. Microbiol. 25;1(9):16116. doi: 10.1038/nmicrobiol.2016.116.

Woese, C., 1998. The universal ancestor. Proc. Natl. Acad. Sci. U S A. 9;95(12):6854–9.

Yang, J., Roy, A., Zhang, Y., 2013. Protein–ligand binding site recognition using complementary binding-specific substructure comparison and sequence profile alignment. Bioinformatics, 29:2588–2595. doi:10.1093/bioinformatics/btt447.

Zahnle, K., Arndt, N., Cockell, C., Halliday, A., Nisbet, E., Selsis, F., Sleep, N.H., 2007. Emergence of a Habitable Planet. Space Sci. Rev. 129:35–78.

Zhang, Y., Skolnick, J., 2005. TM-align: a protein structure alignment algorithm based on the TM-score. Nucleic Acids Res. 22;33(7):2302–9.

Zhang, Y., 2008. I-TASSER server for 3D structure prediction. BMC Bioinformatics, 9:40. doi:10.1186/1471-2105-9-40.

