## Supplementary information for "The structural proteome for the primordial glycolysis/gluconeogenesis"

**Ancestral sequences (Bottom-Up approach)**

>Transketolase
FVQSSLAVGFEPTHSVNSRIFQLQQMASIHLVRPS

>Glucose-6-phosphate-1-dehydrogenase-2
SFSPARGSNPYPEFLSGGPEGLNP

> Glyceraldehyde-3-phosphate-dehydrogenase
CRGFVQSSQGVEPVLARVPERRSGGTQPGPADRR

>Glucose 6P isomerase

PRSFDSFCSGGLGRTRWIEAIVQENTPV

>Glycerate kinase

LICSGHSAVQVSSHELGGRTR

>Triosephosphate isomerase

LSEGVEPVLARVPERLAVGFEPTHSVNS

> Phosphoglycerate kinase

FTLQGPSAVRDVILNRWLRSIWYGPAHHLSGIRS

**Ancestral sequences (Top-down approach- Complete deletion and 99% cutoff)**

>Glycerate kinase

LRTSTYGVGELIRAALHGAERILVGLGGSAQAAQKGARLGIDMVMQDADLVIT
GEGRLTGIALAGI

>Triosephosphate isomerase

LSEGVEPVLARVPERLAVGFEPTHSVNS

>Glucose 6-phosphate 1-dehydrogenase 2

KPFGHDIETVESIETYMAVKPFYIREASWKFVDPILKHYEYESGT

> Glyceraldehyde-3-phosphate-dehydrogenase

VSVVDLTVTYEEEAMEVVSSD

>Transketolase

QPIEQLAMLRSIPNLTVIRPADAAAWLPVLD

**Ancestral sequences (Top-down approach- 95% cutoff)**

>Glyceraldehyde-3-phosphate-dehydrogenase

IRVGINGDIEVVAINDLVDAETMAHLLKYDSVHGRFPGEVEVDDDSLVVNGKK
IKVIAERDPANLPWKELGVDIVMECTGIFTNREKAEAHLEAGAKKVIISAPAKD
YLVNHDQLDPKIISNASCTTNCLAPVAKVLHDKFGGLMTTIILDKDLRRAAQNII
PTSTGAAKAVGLVFPELKGKLNGMSVRVPTPNVSLVDLTFELEKDTTVEEINAV
LKAAAEGILGYTDEPLVSCDYDNEWGFSNRMVDTAVYMAK

>Glucose 6-phosphate1-dehydrogenase

CLVIFGATGDLARRKLIPALYNLYRDGLLPEDFRIIGVARSPWTDEEFREKVREA
LKQHAKAEFDEETWDQFCQRLYYMSGDFDDPDNYAKLAERLEDGNRVFYLST
PPNFFGPICENLAAAGLNGWSRVVIEKPFGHDLESARELNAQLHQVFDENQIYRI
DHYLGKETVQNILVLRFANGIFEPLWNRNYIDHVQITVAETVGVEGRGGYYDS
SGALRDMVQNHLLQLLALVAMEPPASFDADSIRDEKVKVLRSLRPPEDVVRGQ
YTAGTVGGKPVPGYREEPGVSNTETFVAMKLHIDNWRWAGVPFYLRTGKRLP
ERCTEIVIQFKEAPHNIFNDSALSPNRLVIRVQPDEGISLRFNAKTPGAGMRTRQ
VSMDFSYSETFNERSPEAYERLLLDAMRGDQTLFTRSDEVEAAWKFVDPILEH
WEDRGQPHPYPAGSWGPTAADELLARDGRRWHR

>Glucose 6 Phosphate Isomerase (I)

CDIDLEPGTERLMEILWDGPRRLNDMKGILCAFEEMIADIYYMYRDLSPDEDDD
WLHRLRYDTTVIPPNVGGEYAMTKGFHPKAPEGMGYPEILEGEGIMLLQSRRDI
REAEAGDYVFVPNVGPFVMSNLYSTDVHDYYYITEDGLTVVKRWDNAKGPNY
FDEPGIQRYRGSMYGLL

>Glucose 6 Phosphate Isomerase (II)

LKFDYSEELENLEAMEALHNGREEIKEFAQKLREFDNIVVIGIGGSSLGARAVY
GRFFFMNNVDPDSIHELLEHLDKKVAITDFETFPIPDNVGGRFSVLSAGLLPAAI
CGFDIEELLEGARDMNQAFNDDLHQNPVHYRKGKNINVMMPNVGIFPLSAVFT
TDLHSFLQLIQEGDKFVFIKVEKNLDDGKENGRPNMTGYLFGINPFDQPGVEAY
KKNMYGLL

>Glycerate Kinase

MKIVIAPDSFKSLSAEAAAIGPPADGGEGTVAMVAGVGPLGVAGDTAVIEMAA
SGILVRNPTTTGTGELIALDGIIIGIGGSATNDGGAGMALGFLDGGLLILRLVACD
VNPLGGASVFGPQKGAVLDLHAIPGSGAAGGMGGLAFALGEIVLIADLVITGEG
DQTGKPGVAAPVIIAGLGIAFSILAALAI

>Phosphoglycerate kinase

VRVDLNVPLIQDDTRIRAHLPTIKYLQGAVILMAHLGRPKGKFSLAPVAERLSE
LLGRPVKFVDDEVEEAVKNLKPGDVLLLENVRFHPGEEKPDPEFAKQLASLAD
VYVNDAFGTAHRAHASTVGFPFPSAGFLMEKEIEALGKLLDNPERPFVAILGGA
KVSDKIGVIENLLDKVDKLLIGGLMAFTFLKAQGYEIGNSLVEEDKIDLAKELM
EKALLLPVDFVVADDAQTKVVPVDIPILDIGPETVEMFAEALKNAKTVFWNGP
MGVFEMEPFAKGTMAVAQAAESGAVTVVGGGDTAAAVEKFGLADSHISTGG
GASLEFLEGEELPGVEALN

>Triose phosphate isomerase

RKPLIAGNWKMNDAFLLVEVVPLVGIVGAQNDGAFTGEISPLDLGCYVIIGHSE
RRFETDEVNKKALGMTPILCGETLEREAGVQILGLEEVIAYEPVWAIGTGKTAT
AEIRLAEVRILYGGSVKPNEDIDGALVGGASLFI

>Transketolase.

NIQLSTRAASGKVLNELGKKVPNMIVLSADLSNSMKTGDFSNFSGRFIHCGVAE
HNMVGIANGMALHGPFCATFAVFSDYMWPAIRMAALPVKFVWTHDSFSVGED
GPTHQPIEQLAMLRAIPNMHVFYPADATETAAAWKMMLEHKGPTYLVLSRQN
LPVIHKSEEVEKGGILRDDVTLIATGSQVSTALEAAEMLKSVRVVSMPSMELFQ
QQSKKESVLVAVEHMGLHGFVGKVIGMDHFGRSAPAEVLFEKFGFTAENVVQ
KVKE
